# Overexpression of miR-424(322)/-503 Induces Severe Dilated Cardiomyopathy by Regulating The Fatty Acid Oxidation Gene Expression Program

**DOI:** 10.64898/2026.08.17.744731

**Authors:** Shreesti Shrestha, Jiayi Chen, Xiaopeng Shen, Rui Liang, Jahnvi Rajput, Marcello Tosso, Hoang Vu, Anirban Roy, Chin-Yo Lin, Ryan Boudreau, Ashok Kumar, Bradley McConnell, Yu Liu

**Author notes:** These authors contribute equally to this work (Shreesti Shrestha, Jiayi Chen).

## Abstract

Fatty acid oxidation (FAO) is a major energy source in the adult heart, and disruption of cardiac metabolism is closely associated with heart failure. Here, we investigated the effects of cardiac-specific overexpression of the H19X-encoded miR-424(322)/-503 cluster using an inducible transgenic mouse model. Sustained miR-424(322)/-503 overexpression caused rapid metabolic and functional deterioration, with early impairment of fatty acid oxidation. Short-term induction reduced FAO activity and downregulated genes involved in lipid metabolism, while glycolytic enzyme activity remained largely unchanged. Continued miR-424(322)/-503 expression subsequently led to severe dilated cardiomyopathy characterized by ventricular dilation, wall thinning, fibrosis, reduced contractility, and high mortality. Importantly, disease progression was dependent on the level and duration of miR-424(322)/-503 expression, as intermittent or lower-dose induction delayed cardiac dysfunction and prolonged survival. Withdrawal of miR-424(322)/-503 expression after the onset of dysfunction promoted substantial functional recovery. Together, these findings identify miR-424(322)/-503 as a potent regulator of cardiac metabolic reprogramming that disrupts fatty acid metabolism and drives progressive heart failure.

## Introduction

Fatty acids play a central role in maintaining energy balance in the human body, and the heart is especially dependent on them to sustain its high metabolic demands^1^. In a healthy adult heart, mitochondrial fatty acid oxidation (FAO) provides the majority of ATP required for continuous contraction, making efficient FAO essential for normal cardiac function^2^. However, this reliance also means that disruptions in fatty acid metabolism can substantially impair cardiac performance. In heart failure, mitochondrial oxidative capacity often declines, contributing to inadequate ATP production and progressive functional deterioration^3^. This decline in mitochondrial function is now recognized as a major driver of heart-failure pathophysiology. Genetic studies further highlight the importance of intact mitochondrial machinery in maintaining cardiac health. For example, mice lacking Sco2, a gene required for Cytochrome c Oxidase assembly, develop severe cardiomyopathy and myopathy, resulting in early lethality^4^. Mutations in Mitochondrial Trifunctional Protein (MTP), which catalyzes multiple FAO steps, lead to cardiac fibrosis and hepatic steatosis^5^. Similarly, deletion of the mitochondrial fusion genes Mfn1 and Mfn2 causes profound mitochondrial fragmentation; their embryonic loss is lethal, whereas adult deletion rapidly induces dilated cardiomyopathy^5,6^. These findings underscore how sensitive the heart is to mitochondrial structural and metabolic defects.

Clinically, idiopathic dilated cardiomyopathy has been associated with decreased FAO capacity and more rapid depletion of circulating free fatty acids, changes that correlate with reduced cardiac efficiency^7,8^. However, metabolic remodeling in heart failure is heterogeneous. Some studies report reduced myocardial fatty acid uptake and increased glucose utilization, while others find unchanged or even increased fatty acid uptake in certain patient cohorts^9,10^. During early heart-failure development, the heart may temporarily preserve FAO as a compensatory mechanism. For instance, dogs with moderately reduced ejection fraction show no suppression of myocardial FAO^11^. As the disease progresses, however, mitochondrial dysfunction typically limits FAO flux, particularly in conditions such as obesity, metabolic syndrome, or severe pressure-overload–induced heart failure^2^. In Transverse Aortic Constriction (TAC) models, FAO flux and FA-derived ATP production decrease significantly, although fatty acids still remain the predominant source of mitochondrial ATP compared to glucose^12,13^. Together, these observations illustrate that despite metabolic shifts toward greater glucose reliance, FAO continues to play a foundational role in the failing heart^14^.

The H19X locus, located on Xq26.3, encodes several non-coding RNAs, including the miR-424/-503 cluster, miR-542, miR-450a-1/2, and miR-450b, along with a long non-coding host transcript - miR503 host gene^15,16^. miR-322 is the mouse ortholog of human miR-424, thus, we use miR-424(322)/-503 as synonymous to miR- 424/-503. H19X has been identified as a crucial regulator of muscle and cardiac differentiation. Our previous work demonstrated that the H19X-encoded miR-424(322)/-503 cluster promotes cardiomyocyte differentiation by repressing neural-associated pathways through its target Celf1^17^. Notably, miR-424(322)/-503 expression peaks around embryonic day 14 in the developing heart but declines sharply after birth, remaining highly expressed only in the ovary postpuberty. This developmental downregulation suggests a specialized, time-restricted role for H19X in embryonic cardiac differentiation. Despite its postnatal decline, H19X reappears in multiple cardiovascular pathologies. Elevated circulating miR-503 levels have been reported in patients with heart failure with reduced ejection fraction, and known to promote cardiac fibrosis, particularly in the context of angiotensin II signaling^18,19^. Increased miR-424(322) levels have also been linked to fatal acute myocardial infarction and may serve as part of a predictive microRNA panel for future AMI risk, with sex-specific associations^20^. Additional evidence indicates that miR-424(322) contributes to vascular smooth muscle cell dysfunction and interacts with circular RNAs in maintaining vascular homeostasis^21^. Beyond the heart, the H19X-encoded miR-424(322)/-503 cluster also regulates skeletal muscle growth. Its overexpression induces muscle fiber atrophy by targeting translation initiation pathways, while interruption of H19X transcription enhances muscle growth^22^. Elevated miR-424(322) levels have additionally been associated with preoperative skeletal muscle loss, emphasizing its broader role in muscle physiology^23^.

Our study seeks to define the pathological role of H19X by investigating how the H19X-encoded miR-424(322)/-503 cluster regulates cardiac function, fatty acid metabolism, and cardiomyopathic remodeling. By examining the consequences of H19X-encoded miR-424(322)/-503 overexpression in the adult heart, we aim to establish an important non-coding RNA regulator that govern metabolic maturation during maladaptive remodeling in heart disease.

## Methods

### Animal Care and Use

pJG/ Inducible ALPHA MHC was a gift from Jeffrey Robbins (Addgene plasmid # 55792)^24^. A fragment containing the full-length α-MHC promoter, 7 repeats of the TetO sequences, the miR transgene, and the human growth hormone polyadenylation signal was used to develop a mouse strain through Cyagen Biosciece (US).

The pJG-inducible *a*-MHC-miR-424(322)/-503 mice were crossed with homozygous B6. Cg-Gt (ROSA)26Sortm1(rtTA*M2)Jae/J (rtTA) mice (Jackson Laboratory, #006965)^25^ to create the transgenic animals used in this study. The WT and TG mice were fed with a doxycycline diet (Envigo #TD.05512, high dose 2000 mg/kg diet and Envigo #00502, low dose 200 mg/kg diet). All animal studies were approved by the Institutional Animal Care and Use Committee at University of Houston. Mouse strain information is included in Supplement Table 1.

### Cardiac Functional Assessments by Echocardiography

Cardiac function was assessed in mice at multiple time points using a VisualSonics Vevo 3100 high-frequency, high-resolution micro-ultrasound system. Briefly, mice were anesthetized with 1.5 - 2% isoflurane mixed with 1.0 litre/minute 100% O_2_ in the induction chamber. They were then placed on the heating pad with an embedded electrocardiogram (ECG) in a supine position allowing them to receive anesthesia through a nose cone. A toe or tail pinch was done to confirm sedation. The body temperature was continuously monitored through a rectal probe. The heart rates were maintained at 400 – 500 beats/minute (bpm) between experimental groups. The four paws were then taped to ECG electrodes with electrode gel. The chest hair was removed with a hair clipper and ultrasound gel was applied on the chest to image the heart.

Echocardiographic imaging was conducted using a 550D ultrasound probe, with B-mode employed for the long-axis view and M-mode for the short-axis view at the level of the papillary muscles. A standard 2D echocardiographic study was performed in the parasternal short-axis view to assess left ventricular (LV) dimensions and systolic function in B-mode. M-mode echocardiography was used to measure LV posterior wall thickness and interventricular septum thickness at both end-diastole (LVPW,d, IVS,d) and end-systole (LVPW,s, IVS,s), as well as LV internal dimensions at end-diastole (LVID,d) and end-systole (LVID,s). At least three series of three cardiac cycles were recorded per mouse. Image acquisition and subsequent analyses were conducted using Vevo 3100 software, with LV ejection fraction (EF) and LV fractional shortening (FS) calculated by tracing the LV wall.

### Tissue Harvesting and Processing

Mice were euthanized and the hearts were removed by dissecting the mice to open the chest cavity. The hearts were cut into two pieces; one was quickly snapped frozen and stored at -80°C for RNA while the other piece was immersed into 4% PFA for overnight fixation at 4°C. The next day, hearts were rinsed twice in PBS for 10 minutes each, and once in 70% ethanol. They were then stored in 70% ethanol until further processing.

For paraffin-embedding, the hearts were processed in a series of increasing alcohol concentrations (70%, 85%, and 100%) for an hour each. The dehydrated hearts were then embedded in two changes of xylene for an hour and three changes of paraffin. The processed hearts were finally embedded using paraffin in either longitudinal or transverse sections depending on downstream analysis.

### Histology

Formalin-fixed, paraffin-embedded heart tissues were sectioned at 5 μm using a microtome. Tissue sections were deparaffinized in xylene, followed by a graded ethanol series for rehydration. H&E staining was performed using a Sigma-Aldrich kit. The deparaffinized sections were stained in haematoxylin for 10 minutes followed by rinsing. The sections were then decolorized using 0.1% acid alcohol. Eosin Y stain was applied for 30 seconds followed by immediate 95% ethanol wash.

Fibrosis was assessed using a Masson’s trichrome staining kit (Sigma Aldrich, St. Louis, MO, USA). Tissue sections were first preheated in bouin’s solution at 56°C for 15 minutes. After cooling down to RT, sections were incubated in Weigert’s iron hematoxylin for 5 minutes to stain nuclei black. This was followed by staining with Biebrich scarlet-acid fuchsin for 5 minutes, which marked acidophilic structures such as cytoplasm, muscle, and collagen in red. Subsequently, aniline blue was applied to stain collagen fibers blue, enabling the differentiation of fibrotic regions. Finally, sections were rinsed in 1% acetic acid.

Collagen deposition was further analyzed using a Picro-Sirius Red Staining Kit (Abcam, Cambridge, UK). Deparaffinized sections were pretreated with 0.2% phosphomolybdic acid for 5 minutes, followed by incubation in Picro-Sirius Red solution for 1 hour to specifically stain collagen fibers. Slides were then subjected to 2 washes with 0.5% acetic acid.

### RNA Isolation and Quantitative Real-Time PCR

Total RNA was isolated using Ribozol reagent (Genedepot) according to the manufacturer’s protocol with the following modification. After transferring the aqueous phase to a 1.5 ml eppendorf tube, absolute ethanol was added and placed at -20°C overnight for the precipitation phase. For assessing the miRNA expression, cDNA synthesis and qRT-PCR were set up using TaqMan miRNA Reverse Transcription Kit (ThermoFisher Scientific) and Takyon^TM^ One-Step Kit Converter (Eurogentec) according to the manufacturer’s instructions. First, 2 ng of total RNA was reversed transcribed using miRNA stem-loop primers: mmu-miR-424 (#001076, ThermoFisher Scientific) and mmu-miR-503 (#002456, ThermoFisher Scientific). qRT-PCR was then run with Takyon^TM^ One-Step Kit Converter (Eurogentec). miRNA expression was normalized to 18s RNA. qRT-PCR was performed (Applied Biosystems 7500) using the cycling conditions: 48°C for 30 minutes (Reverse transcription step); 95°C for 10 minutes (Initial denaturation); 40 cycles of 95°C for 15 seconds and 60 °C for 60 seconds (Extension).

Quantitative real-time PCR (qRT-PCR) (Applied biosystems 7500) for mRNA genes was done using Takyon ROX SYBR MasterMix dTTP (Eurogentec) following the manufacturer’s instructions. 100 ng of total RNA was used as the starting concentration. GAPDH was used as endogenous control. The PCR primers used for qRT-PCR are included in Supplement Table 2. The cycling condition used were: 48°C for 10 minutes (Reverse transcription step); 95°C for 3 minutes (Initial denaturation); 40 cycles of 95°C for 10 seconds, 60°C for 60 seconds, and 72°C for 30 seconds.

### Assessment of FAO and Glycolytic Enzyme Activities

Frozen or freeze-thawed tissues were mechanically homogenized in ice-cold 1X lysis or sample buffer, centrifuged at ∼14,000 rpm for 5 min at 4°C, and supernatants were collected for enzymatic assays or stored at −80°C. Protein concentrations were determined using the bicinchoninic acid (BCA) assay. Fatty acid oxidation (FAO) using BMRservice #E-141 and hexokinase (HK) using BMRservice #E-111 activities were measured using colorimetric assays in 96-well plates, with samples assayed in duplicate. Reaction mixtures containing either control solution or substrate solution were incubated at 37°C for 60 or 120 min in a non-CO₂ humidified incubator, and absorbance was measured at 492 nm. Enzyme activities were calculated from the change in optical density (ΔO.D.) using manufacturer-provided conversion factors and normalized to protein concentration. Pyruvate kinase (PK) using BMRservice #E-117 activity was measured using a coupled ATP luminescence assay, in which reactions were initiated by substrate addition, incubated at 37°C for 1 min, terminated with trichloroacetic acid, clarified by centrifugation, and quantified by luminometry. PK activity was calculated as ΔRLU relative to control reactions and normalized to protein concentration.

### RNA Sequencing Analysis

Frozen heart tissues from both wild-type (WT) and transgenic (TG) mice, which were fed doxycycline for 14 days, were thawed prior to RNA extraction. RNA isolation was performed using Trizol reagent, following the protocol outlined earlier. Triplicate samples from each group were submitted for RNA sequencing at Novogene. A list of differentially expressed genes (DEGs) was obtained from Novogene. The up-regulated and down-regulated genes were identified and separated using DESeq2 package in R^27^. Gene set enrichment analysis was conducted using GSEA 4.1^28,29^ and ShinyGo 0.80^30^ online tools. The lists of up-regulated and down-regulated genes were uploaded to the database for annotation, visualization, and integrated discovery (DAVID). The heatmap of WT and TG samples from mice was generated using the list of differentially expressed genes with log_2_ fold change > 1 and padj < 0.01.

### Statistical Analysis

The results are given as mean ±SEM. All statistical analyses were performed using GraphPad Prism Software (Version 9.0). Unpaired 2-tailed Student t-tests were used to determine the significance between 2 groups, considering p < 0.05 as statistically significant. Analyses between multiple groups were performed using 1-way ANOVA or 2-way ANOVA following Tukey multiple comparison posttest to determine significance.

Echocardiography results were analyzed by 2-way ANOVA or by 2-way ANOVA repeated measures test when the same animal was used over the course of the experiments. P < 0.05 was considered statistically significant.

## Results

### miR-424(322)/-503 Overexpression Induces Rapid Dilated Cardiomyopathy

To study the function of miR-424(322)/-503 in adult cardiomyocytes, we created a miR-424(322)/-503 transgenic mouse strain (TG) in which miR-424(322)/-503 is induced in cardiomyocytes by a doxycycline-containing chow (2000 mg/kg diet). 8–10-week-old wild-type (WT) and transgenic (TG) mice were placed on doxycycline chow for 28 days. Cardiac function was assessed by echocardiography two days before doxycycline administration to establish baseline measurements and subsequently monitored every 7 days (Figure 1A). By day 28, miR-424(322) and miR-503 expression levels were elevated in TG hearts (Figure 1B). TG mice developed significantly enlarged hearts compared to WT controls, with an increased heart-weight-to-body-weight ratio (Figure 1C). TG mice also experienced a progressive decline in body weight, in contrast to WT mice maintained on a doxycycline diet (Figure 1F). By the study endpoint, TG mice were visibly thinner than WT animals, consistent with sustained systemic wasting (Figure 1E). Survival analysis showed that ∼60% of TG mice died within the 28-day induction period, indicating the severe physiological burden caused by miR-424(322)/-503 overexpression (Figure 1D). Together, these findings indicate that forced miR-424(322)/-503 expression triggers marked cardiac remodeling accompanied by profound systemic deterioration.

**Figure 1.**
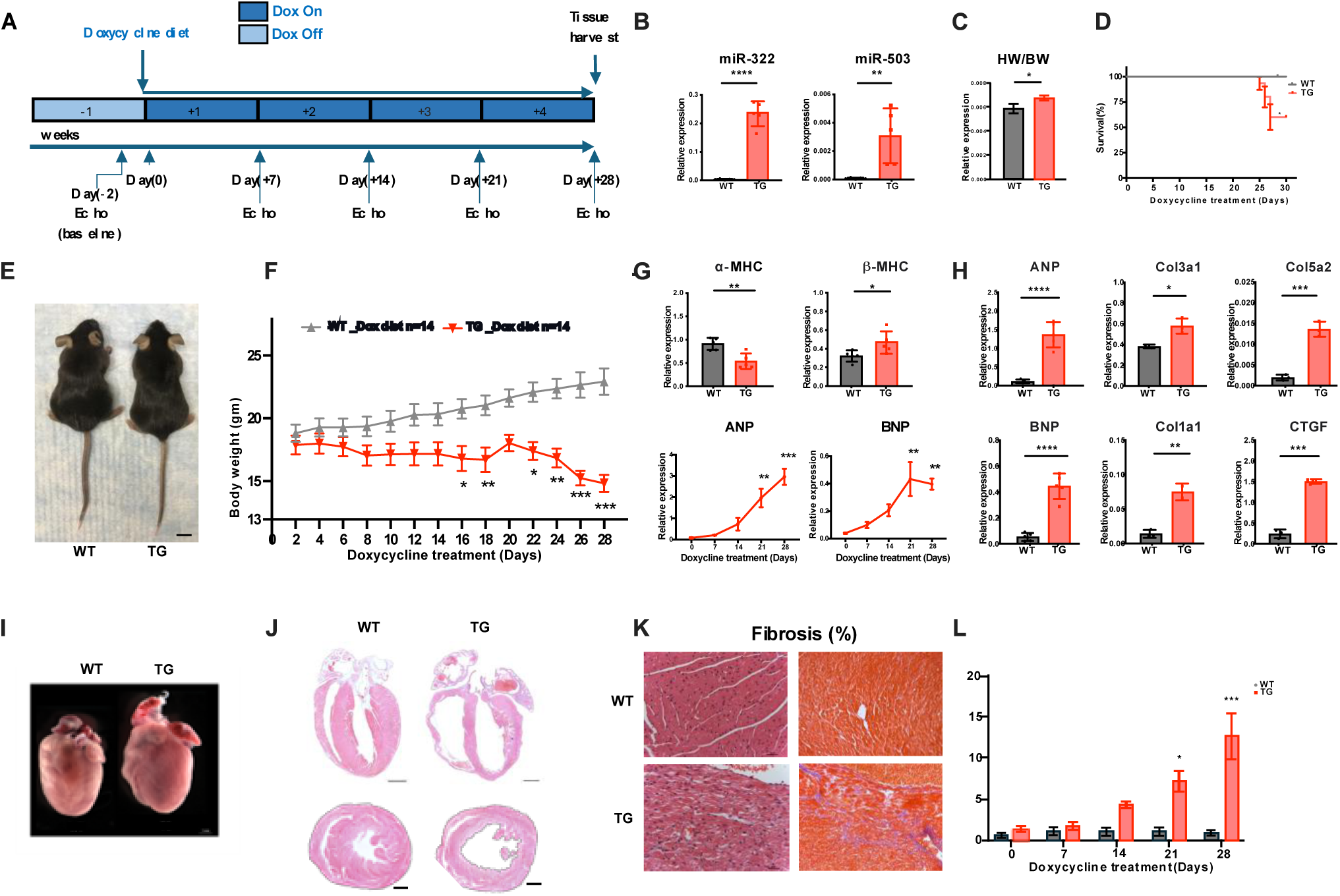
H19X-encoded miR-424(322)/-503 overexpression led to heart failure in mice. **(A)** Schematic timeline of doxycycline induction. Wild-type (WT) and transgenic (TG) mice were fed a doxycycline-containing diet to induce miR-424(322)/-503 expression. Baseline cardiac function was assessed 2 days prior to induction, followed by serial measurements every 7 days through day 28. **(B)** qPCR analysis of miR-424(322) and miR-503 expression levels in WT and TG hearts at day 28 (WT n=8, TG n=5, student’s t-test). **(C)** Heart-to-body weight ratios of WT and TG mice at day 28 (WT n=8, TG n=5, student’s t-test). **(D)** Kaplan–Meier survival analysis over a 30-day period (WT n=10, TG n=15, log-rank test). **(E)** Representative whole-body images of WT and TG mice at day 28. Scale bar, 1 inch. **(F)** Body weight measurements of WT and TG mice maintained on doxycycline diets (WT n=14, TG n=14, two-way ANOVA). **(G)** qPCR analysis of cardiac hypertrophic markers α-MHC and β-MHC at day 28 (WT n=5, TG n=5, student’s t-test), and cardiac stress markers ANP and BNP over the 28-day time course (TG n=3-5, one-way ANOVA). **(H)** qPCR analysis of cardiac stress markers ANP and BNP, and fibrosis-associated genes Col3a1, Col5a2, Col1a1, and CTGF at day 28 (WT n=5, TG n=5, student’s t-test). **(I)** Representative gross morphology of whole hearts from WT and TG mice at day 28. **(J)** Hematoxylin and eosin (H&E) staining of WT and TG hearts in longitudinal and transverse sections. Scale bar, 500 μm. **(K)** Higher-magnification images of H&E-stained WT and TG heart sections (left) and corresponding Masson’s trichrome staining (right). **(L)** Quantification of myocardial fibrosis based on picrosirius red staining at days 0, 7, 14, 21, and 28 (WT n=3, TG n=3, two-way ANOVA). *Date is represented as mean ± SEM. * p < 0.05*, *\*\* p < 0.01; *** p < 0.001 **** p < 0.0001*.

TG mice exhibited molecular signatures of heart failure by day 28. A switch from α-MHC to β-MHC was evident (Figure 1G, top), reflecting a transition from adult to fetal contractile isoforms. The fetal gene program was progressively reactivated, with ANP and BNP levels rising over the induction period, while α-MHC declined significantly as early as day 14 (Figure 1G bottom; Supplement Figure 3C). At day 28, ANP and BNP were drastically elevated and collagen-related genes (Col1a1, Col3a1, Col5a2, CTGF) were markedly upregulated in TG hearts as well (Figure 1H).

Histological assessment further demonstrated dilated cardiomyopathy–like remodeling. H&E staining revealed left ventricular thinning and chamber dilation in both longitudinal and transverse sections of TG hearts (Figure 1J). High-magnification views showed myocyte disorganization, and Masson’s trichrome staining identified interstitial fibrosis (Figure 1K). To evaluate disease progression, hearts were collected at days 0, 7, 14, 21, and 28. By day 14 TG hearts displayed emerging chamber dilation and wall thinning that worsened through day 28 (Supplement Figure 3A top). Picrosirius red staining confirmed fibrosis beginning at day 14, increasing by approximately 13% by day 28 (Figure 1L, Supplement Figure 3A bottom).

Echocardiography showed progressive deterioration of systolic pump function (Figure 2). Both WT and TG mice exhibited normal ejection fraction (EF) and fractional shortening (FS) values at baseline (day 0) prior to miR-424(322)/-503 induction. Following miR-424(322)/-503 induction, TG mice displayed a gradual decline in EF and FS, starting at day 14 of induction (Figure 2B). EF dropped from 60.66 ± 5.98% at day 0 to 15.35 ± 9.70% at day 28, while FS decreased from 32.06 ± 4.55% to 6.96 ± 4.64% over the same period (Table 1). Cardiac output in TG mice declined from 16.68 ±2.41 ml/min to 5.26 ± 2.40 ml/min, and stroke volume dropped from 37.92 ± 6.07 μl to 13.20 ± 7.67 μl, further demonstrating impaired cardiac function (Table 1, Figure 2C). Additionally, left ventricular internal diameter during diastole (LVID,d) and systole (LVID,s) increased significantly in TG mice, indicating ventricular chamber dilation (Figure 2D). Conversely, interventricular septum thickness during diastole (IVS,d) and systole (IVS,s), as well as left ventricular posterior wall thickness during diastole (LVPW,d) and systole (LVPW,s), decreased markedly, reflecting significant thinning of the ventricular walls (Figure 2E,F). Collectively, these findings indicate that miR-424(322)/-503 overexpression significantly impairs cardiac function, leading to ventricular dilation, reduced wall thickness, and decreased contractility which are hallmark features of dilated cardiomyopathy^31^. In past studies, such a rapid and severe disease phenotype was often caused by ablating genes for sarcomere proteins, nuclear envelop proteins, cytoskeletal proteins or combing stress signaling and chemotherapy drugs. Generic manipulation of other miRNAs rarely produce such a rapid and severe cardiac function deterioration^32^. To evaluate potential sex-based differences, we compared echocardiographic measurements in male and female mice. Following 28 days of miR-424(322)/-503 induction, EF decreased from 61.09 ± 6.58% to 15.27 ± 10.84% in males and from 57.88 ± 5.87% to 23.48 ± 5.02% in females (Table 2). Similarly, FS declined from 32.41 ± 5% to 6.94 ± 5.18% in males and from 29.8 ± 3.97% to 10.7 ± 2.47% in females. Both sexes exhibited increased left ventricular internal diameter (LVID) alongside decreased interventricular septum thickness (IVS) and left ventricular posterior wall thickness (LVPW). These findings indicate that cardiac function was significantly impaired in both males and females, suggesting that miR-424(322)/-503 overexpression induces heart failure irrespective of sex. Further, females appeared to retain more systolic function than males, agreeing with the frequent observation that females are more resistant to cardiac insults in rodent models and humans^33–39^.

**Figure 2.**
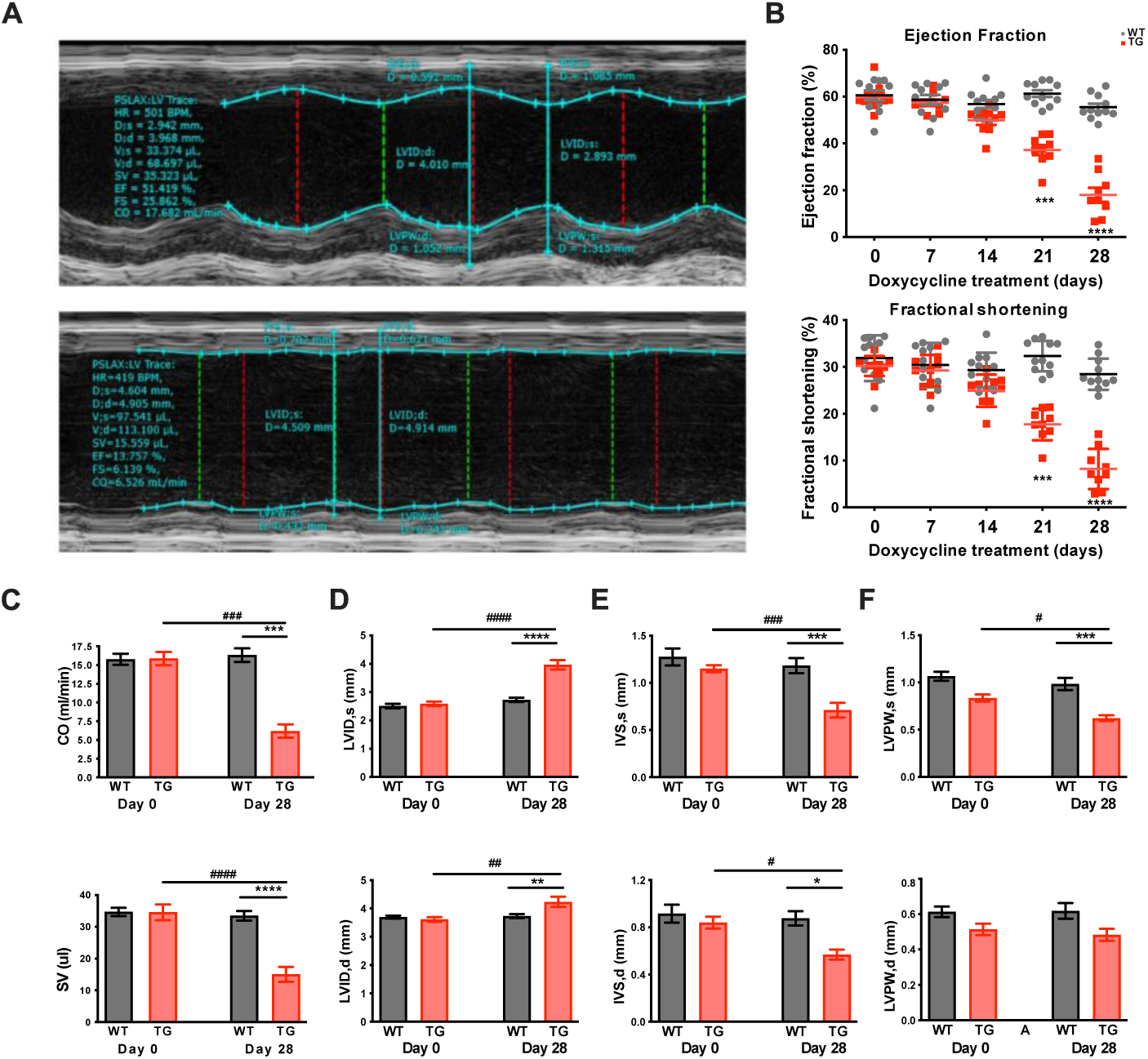
H19X-encoded miR-424(322)/-503 overexpression causes progressive cardiac dysfunction. **(A)** Representative echocardiographic images illustrating systolic and diastolic cardiac parameters in WT and TG mice. **(B)** Longitudinal echocardiographic assessment of left ventricular systolic function, including ejection fraction (EF, %) and fractional shortening (FS, %), measured at days 0, 7, 14, 21, and 28 following miR-424(322)/-503 induction (WT n=11, TG n=10, two-way ANOVA). **(C)** Quantitative analysis of cardiac output (CO, mL/min) and stroke volume (SV, μL) in WT and TG mice at baseline (day 0) and day 28 (WT n=11, TG n=9, two-way ANOVA). **(D)** Quantitative analysis of left ventricular internal diameter at systole (LVID;s) and diastole (LVID;d) at baseline and day 28 (WT n=11, TG n=9, two-way ANOVA). **(E)** Quantitative analysis of interventricular septal thickness at systole (IVS;s) and diastole (IVS;d) at baseline and day 28 (WT n=11, TG n=9, two-way ANOVA). **(F)** Quantitative analysis of left ventricular posterior wall thickness at systole (LVPW;s) and diastole (LVPW;d) at baseline and day 28 (WT n=11, TG n=9, two-way ANOVA). *Date is represented as mean ± SEM. * p < 0.05*, *\*\* p < 0.01; *** p < 0.001 **** p < 0.0001 vs Day 28 WT; * p < 0.05*, *\*\* p < 0.01; *** p < 0.001 **** p < 0.0001 vs Day 0 TG*.

### Intermittent or Low-Level miR-424(322)/-503 Expression Delays Heart Failure Progression

To investigate whether the progression of miR-424(322)/-503-induced heart failure could be attenuated by reducing miR-424(322)/-503 expression, we administered an intermittent doxycycline protocol (2000 mg/kg diet). The treatment protocol consisted of three days on a doxycycline diet (dox on) followed by four days on a control diet (dox off) continued for 8 weeks (Supplement Figure 1A). Cardiac function was assessed weekly, and tissue samples were collected at week 8. Intermittent doxycycline administration resulted in a lower expression of miR-424(322) and miR-503 compared to continuous doxycycline exposure (Supplement Figure 1B). α-MHC expression was downregulated, while β-MHC expression was upregulated by week 8 (Supplement Figure 1C top). Additionally, cardiac failure markers, ANP and BNP, were significantly elevated in TG mice (Supplement Figure 1C bottom), suggesting that intermittent doxycycline treatment mitigates the rate of heart failure progression compared to continuous administration.

Notably, in contrast to the continuous treatment group, where TG mice exhibited severe heart failure and mortality within one month of miR-424(322)/-503 induction, none of the TG mice subjected to the intermittent protocol died within this period. Instead, survival was extended to at least two months with intermittent miR-424(322)/-503 levels. Echocardiographic analysis revealed a significant decline in cardiac function from week 5, as indicated by a progressive reduction in ejection fraction (EF) and fractional shortening (FS) (Supplement Figure 1G). Specifically, EF decreased from a baseline of 63 ±5% to 29 ± 10% by week 8, while FS declined from 33.46 ± 3.3% to 15 ± 5.4%. Furthermore, LVID,s and LVID,d exhibited an increasing trend (Supplement Figure 1D), whereas IVS,d and IVS,s, with LVPW,s and LVPW,d decreased over the 8-week intermittent treatment period (Supplement Figure 1E,F).

To further understand the dose-dependent effects of miR-424(322)/-503 on cardiac function, we continuously administered doxycycline at a tenfold lower concentration (200 mg/kg diet). Under this regimen, TG mice exhibited an even greater extension in survival compared to both the intermittent and continuous high-dose doxycycline groups. Cardiac function declined gradually over time, as reflected by a progressive decrease in EF and FS, but at a slower rate than that observed with intermittent doxycycline treatment (Supplement Figure 3B).

### Withdrawal Of miR-424(322)/-503 Expression Reverses Heart Failure

To further investigate the causal role of miR-424(322)/-503 in heart failure progression, its expression was interrupted following the onset of cardiac dysfunction. miR-424(322)/-503 was continuously induced for 14 days, after which the doxycycline-enriched chow (2000 mg/kg diet) was replaced with standard chow to cease further induction (Figure 3A). The mice were allowed to recover, and cardiac tissues were collected at week 12. By the study’s endpoint, the expression levels of both miR-424(322) and miR-503 had returned to near-baseline levels (Figure 3B). ANP and BNP levels increased, while α-MHC expression decreased within the first 14 days of miR-424(322)/-503 induction. However, by week 12, ANP and BNP levels had significantly decreased, while α-MHC expression was restored following 10 weeks of doxycycline withdrawal, suggesting a reversal of molecular markers associated with heart failure (Figure 3C, Supplement Figure 3E).

**Figure 3.**
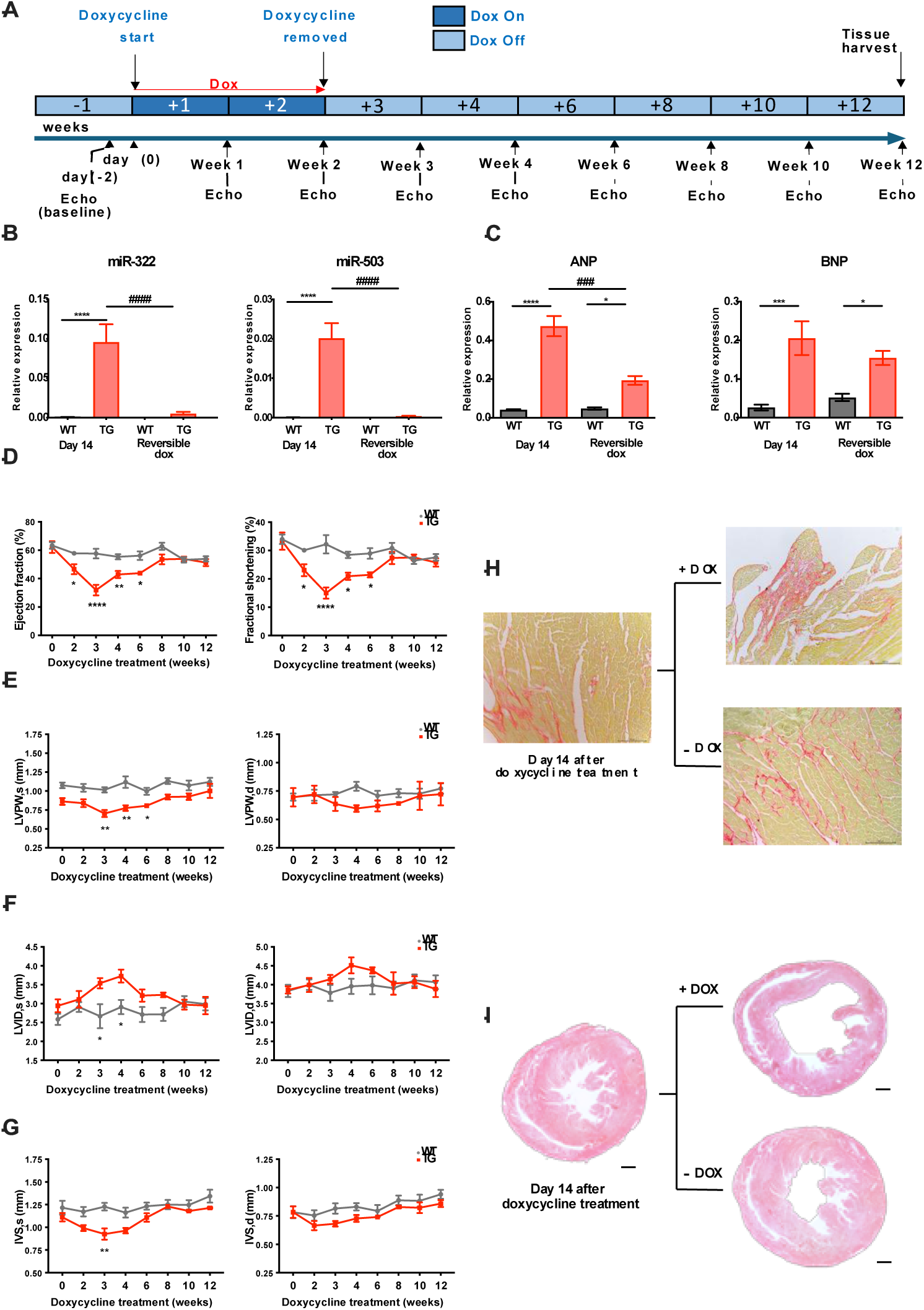
Withdrawal of miR-424(322)/-503 expression promotes functional recovery and reverses cardiac remodeling. **(A)** Schematic timeline of doxycycline withdrawal. Wildtype (WT) and Transgenic (TG) mice were fed a doxycycline-containing diet for 2 weeks to induce miR-424(322)/-503 expression (Dox on), followed by doxycycline withdrawal for 10 weeks (Dox off). Baseline cardiac function was assessed 2 days prior to induction, with serial echocardiographic measurements performed weekly through week 12. **(B)** qPCR analysis of miR-322 and miR-503 expression levels at day 14 (week 2) and week 12 following doxycycline withdrawal (WT n=6, TG n=5, two-way ANOVA). **(C)** qPCR analysis of cardiac stress markers ANP and BNP at day 14 (week 2) and week 12 (WT n=6, TG n=5, two-way ANOVA). **(D)** Longitudinal echocardiographic assessment of left ventricular systolic function, including ejection fraction (EF, %) and fractional shortening (FS, %), measured at weeks 0, 2, 3, 4, 6, 8, 10, and 12 following miR-424(322)/-503 withdrawal on week 2 (WT n=6, TG n=5, two-way ANOVA). **(E)** Quantitative analysis of left ventricular posterior wall thickness at systole (LVPW;s) and diastole (LVPW;d) over the same time course (WT n=6, TG n=5, two-way ANOVA). **(F)** Quantitative analysis of left ventricular internal diameter at systole (LVID;s) and diastole (LVID;d) over the same time course (WT n=6, TG n=5, two-way ANOVA). **(G)** Quantitative analysis of interventricular septal thickness at systole (IVS;s) and diastole (IVS;d) over the same time course (WT n=6, TG n=5, two-way ANOVA). **(H)** Representative Masson’s trichrome staining of TG heart sections with continued doxycycline exposure or following doxycycline withdrawal after 14 days of miR-424(322)/-503 induction. Scale bar, 50 μm. **(I)** Representative hematoxylin and eosin (H&E) staining of TG transverse heart sections under continued doxycycline treatment or after doxycycline withdrawal. Scale bar, 500 μm. *Date is represented as mean ± SEM. * p < 0.05*, *\*\* p < 0.01; *** p < 0.001 **** p < 0.0001 vs Day 14 WT; * p < 0.05*, *\*\* p < 0.01; *** p < 0.001 **** p < 0.0001 vs Day 14 TG*.

Cardiac function was assessed weekly using echocardiography. While WT mice maintained normal EF and FS values throughout the study, TG mice exhibited a decline in EF from a baseline of 62.25 ± 8.3% (week 0) to 47.1 ± 6.6% at week 2. Due to the time required for doxycycline clearance, EF continued to decline, reaching 32± 7.55% by week 3. However, once miR-424(322) and miR-503 expression ceased, cardiac function showed a progressive recovery, as indicated by an increase in EF from 31.92 ± 7.55% at week 3 to 51.27 ± 5.02% by week 12. Similarly, FS declined from 33.27± 6.07% at baseline to 23.06 ± 4.23% at week 2 and 14.98 ± 3.95% at week 3, followed by an increase to 25.79 ± 2.92% upon miR-424(322)/-503 withdrawal (Figure 3D). Structural changes in the heart, including increased LVID and reduced LVPW and IVS thickness during miR-424(322)/-503 induction, also returned to near-normal levels following the cessation of miR-424(322)/-503 expression (Figure 3E,F,G). These findings indicate that continuous miR-424(322)/-503 expression is essential for heart failure progression, and its removal leads to functional recovery.

Histological analysis further supported these findings. H&E and Picrosirius Red staining revealed that myocardial dilation and fibrosis did not worsen over time after the cessation of miR-424(322)/-503 induction (Figure 3H,I). This suggests that upon withdrawal of exogenous miR-424(322)/-503 expression, cardiac function can recover to near-normal levels without regression of the mild chamber dilation and fibrotic remodeling. In other studies, removing various cardiac insults often led to incomplete regression of fibrosis^40,41,42,43^. Collectively, these results demonstrate that miR-424(322)/-503 is required for heart failure progression and that miR-424(322)/-503-induced cardiac dysfunction is reversible.

### Transcriptomic Profiling Reveals Extensive Metabolic and Structural Remodeling

To characterize transcriptomic alterations associated with miR-424(322)/-503 overexpression, we performed genome-wide RNA-seq analysis on WT and TG hearts harvested at day 14 of doxycycline induction. This timepoint was selected based on echocardiographic evidence showing the earliest detectable decline in cardiac function. Sequencing generated reads from 35,735 genes, of which 13,926 genes displayed non-zero counts.

Principal component analysis (PCA) revealed tight clustering of biological replicates within each genotype, confirming dataset consistency (Figure 4A, top). WT and TG samples segregated distinctly along PC1, which accounted for ∼80% of total variance, indicating widespread and almost uniform transcriptional reprogramming induced by miR-424(322)/-503. Differential expression analysis identified 4,356 DEGs between WT and TG hearts. Among these, 1,893 genes were significantly upregulated (padj < 0.05) and 2,463 were significantly downregulated (Figure 4A, bottom). Heatmap visualization further demonstrated clear genotype-specific expression patterns (Figure 4A, right). Cumulative fraction analysis (CFA) revealed a leftward shift of predicted miR-424(322)/-503 target genes in transgenic hearts, indicating preferential downregulation consistent with target-specific repression (Supplement Figure 4). In contrast, predicted targets of other abundant cardiomyocyte miRNAs showed no significant shifts, suggesting that miR-424(322)/-503 overexpression does not globally disrupt miRNA-mediated regulation.

**Figure 4.**
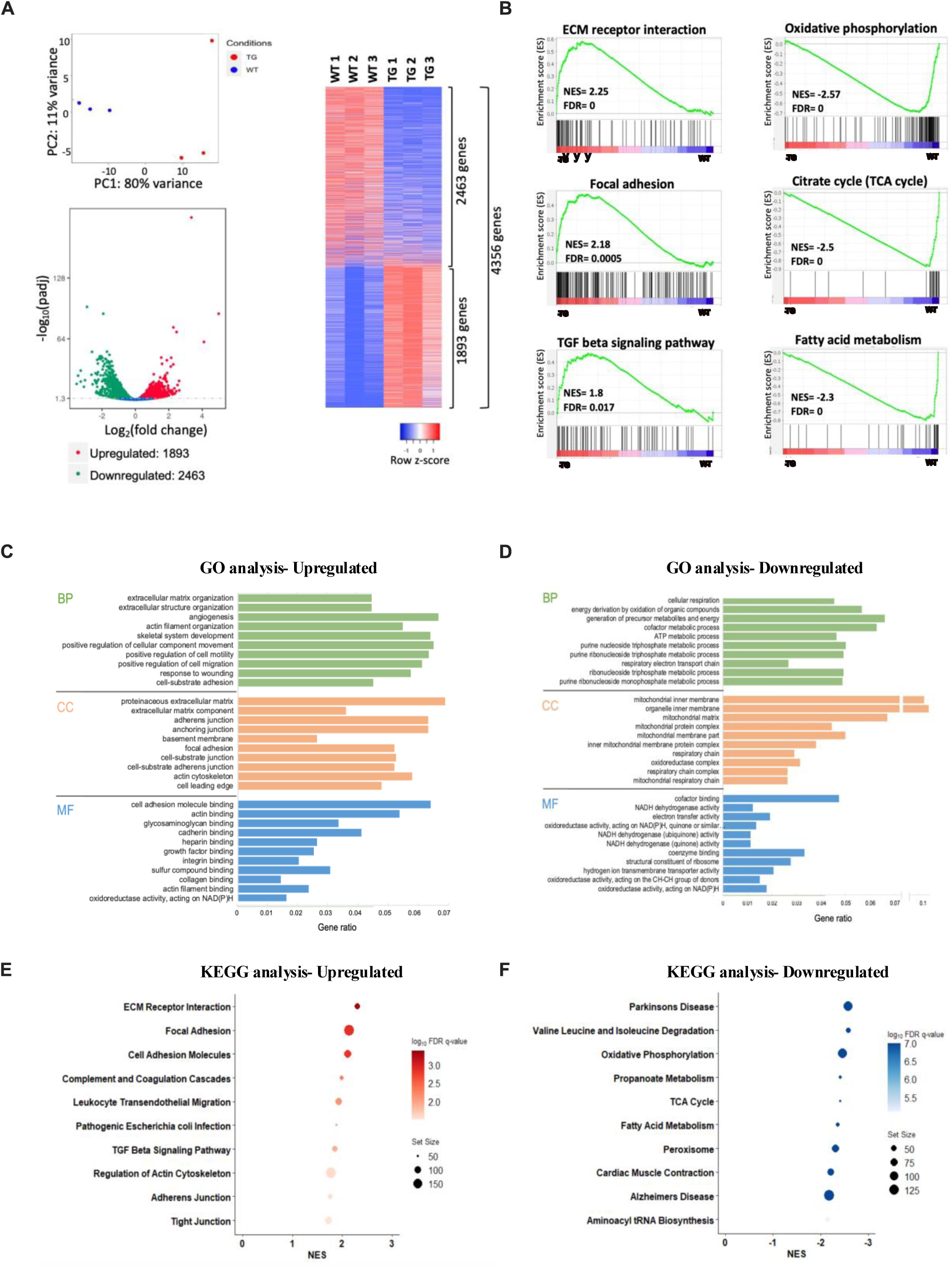
Transcriptomic profiling reveals activation of cardiac remodeling pathways and suppression of metabolic programs in TG hearts. **(A)** Principal component analysis (PCA; top left) demonstrating separation between wild-type (WT) and transgenic (TG) cardiac transcriptomes, volcano plot (bottom left) showing differentially expressed genes, and heatmap (right) of 4,356 differentially expressed genes (DEGs), including 1,893 significantly upregulated and 2,463 significantly downregulated genes in TG hearts. **(B)** Gene Set Enrichment Analysis (GSEA) of transcriptomic changes in TG hearts, highlighting enrichment of extracellular matrix (ECM)–related and pro-fibrotic pathways (ECM–receptor interaction, focal adhesion, TGF-β signaling) and suppression of metabolic pathways including oxidative phosphorylation, tricarboxylic acid (TCA) cycle, and fatty acid metabolism. **(C-D)** Gene Ontology (GO) enrichment analysis of the top 10 biological process (BP), cellular component (CC), and molecular function (MF) categories among upregulated and downregulated genes. **(E-F)** Kyoto Encyclopedia of Genes and Genomes (KEGG) pathway enrichment analysis of significantly upregulated and downregulated pathways in TG hearts.

The top 25 upregulated and downregulated genes are listed (Table 6). Gene Set Enrichment Analysis (GSEA) revealed positive enrichment for extracellular matrix (ECM) receptor interaction, focal adhesion, and TGF-β signaling pathways, confirming robust activation of structural remodeling programs in TG hearts. Conversely, gene sets related to oxidative phosphorylation, the TCA cycle, and fatty acid metabolism displayed negative enrichment scores, indicative of diminished mitochondrial and metabolic function. (Figure 4B)

Gene Ontology (GO) and KEGG pathway analyses of significantly upregulated and downregulated genes further supported these findings (Figure 4C–F). For biological processes (BP), upregulated genes were enriched in ECM organization, angiogenesis, and cell migration. In the cellular component (CC) category, upregulated genes mapped to focal adhesion sites, ECM components, and the actin cytoskeleton. Regarding molecular function (MF), upregulated genes were enriched for adhesion molecule binding, actin binding, and collagen binding. (Figure 4C) For downregulated genes, BP downregulated genes were associated with cellular respiration, electron transport chain activity, and ATP production. In CC, downregulated genes localized to the mitochondrial inner membrane, protein complexes, and respiratory chain machinery. In MF, downregulated genes involved NADH dehydrogenase activity, electron transfer, and oxidoreductase activity. (Figure 4D) KEGG pathway analysis further confirmed GO analysis. (Figure 4E,F) Collectively, these results indicate that miR-424(322)/-503 drives robust activation of ECM and cytoskeletal remodeling programs while suppressing mitochondrial gene networks central to oxidative metabolism.

To assess dynamic changes during disease progression, RNA-seq datasets from TG hearts at day 0 (D0) and day 14 (D14) were compared. Consistent with WT–TG comparisons, GSEA, GO, and KEGG analyses identified substantial pathway remodeling between D0 and D14. Upregulated pathways prominently included PI3K–AKT signaling and several overlapping remodeling-associated pathways. (Supplement Figure 2A). Downregulated pathways were dominated by mitochondrial processes, including oxidative phosphorylation, the TCA cycle, β-oxidation, and fatty acid metabolism, highlighting impaired ATP-generating capacity (Supplement Figure 2B).

In D14 TG hearts, upregulated BP terms reflected enhanced ECM organization, cell adhesion, and TGF-β signaling. CC terms were enriched for extracellular structures, cell-surface compartments, and cytoskeletal elements, while MF terms included ECM binding and protein interaction functions (Supplement Figure 2C). In contrast, downregulated BP terms included fatty acid metabolism, lipid oxidation, and mitochondrial energy production. Downregulated CC terms were dominated by mitochondrial localization, including inner membrane and respiratory chain components, and MF terms comprised NADH dehydrogenase and oxidoreductase activities (Supplement Figure 2D). Gene Set Enrichment Analysis (GSEA) was preformed to further confirm the biological significance of both the upregulated and downregulated genes. In downregulated genes, these were significantly enriched in pathways related to oxidative phosphorylation, fatty acid metabolism, and adipogenesis, suggesting a reduction in energy production and metabolic efficiency in the D14 hearts. In upregulated genes, these were predominantly involved in apoptosis-related pathways, such as the apoptosis pathway, p53 pathway, and TNFα signaling pathway and remodeling pathway like epithelial-to-mesenchymal (EMT) transition. (Supplement Figure 2F)

Additionally, a volcano plot was generated to visually represent the differentially expressed genes (DEGs) between D0 and D14. Specifically, genes related to fatty acid oxidation (FAO) were highlighted in green, revealing many of the FAO-related genes were downregulated. (Supplement Figure 2E) The most significantly downregulated FAO-related genes were labeled in the plot, and a list of these genes was also compiled for further analysis (Table 7). Together, these transcriptomic analyses demonstrate that miR-424(322)/-503 overexpression initiates early and profound suppression of mitochondrial metabolic pathways while simultaneously activating ECM, cytoskeletal, and TGF-β–mediated remodeling programs, providing a molecular basis for the structural and functional deterioration observed in TG hearts.

### Short-Term miR-424(322)/-503 Overexpression Impairs Fatty Acid Metabolism

To exclude the possibility that the change in FAO-related gene expression is secondary to heart failure, instead of being a primary function of miR-424(322)/-503, we analyzed mice with 7 days of miR-424(322)/-503 induction. These mice showed normal cardiac function in echocardiography.

TG hearts exhibited a marked reduction in FAO activity, whereas glycolytic activity as indicated by hexokinase and pyruvate kinase assays, remained unchanged (Figure 5A). A schematic overview of regulatory relationships within fatty acid metabolic pathways is shown in Figure 5B. Three enzymes ATGL, HSL, MGL and a mandatory activator CGI-58 are required in lipolysis^44^. While HSL and CGI-58 levels remained unchanged, ATGL and MGL expression levels were significantly reduced in TG hearts, suggesting compromised lipolytic activity (Figure 5C). Critical FAO regulator, MLYCD, degrades malonyl CoA which is a potent inhibitor of mitochondria fatty acid uptake^45,46,47^, CPT2 regenerates fatty acyl-CoA, making ready for β oxidation^48,49^. Both were significantly downregulated, implying impaired mitochondrial fatty acid transport (Figure 5C). Expression of the lipid droplet–associated genes FITM1 and FITM2 was significantly decreased, indicating reduced cardiac lipid use^50^. Interestingly, the upstream transcriptional regulators PPARGC1α and PPARα were upregulated, likely reflecting a compensatory response to diminished FAO capacity (Figure 5E). In parallel, ACSL1, the enzyme responsible for activating long-chain fatty acids for either β oxidation or synthesis of critical structural lipids^51–54^, was significantly decreased. ACADM and other core enzymes in the four-step β-oxidation remained unchanged, suggesting that downstream mitochondrial oxidation was not directly affected. ARL2 is one of the most downregulated genes in the TG transcriptome. It is a small GTPase functioning primarily in regulating microtubule dynamics but also known to enhance mitochondrial fusion^55,56^. ARL3, a close cousin without mitochondrial function, showed no changes (Figure 5F). This agrees with earlier studies that early heart failure is associated with increased PPARGC1α and PPARα whereas late stage heart failure indicated by decline in systolic function almost always featured a reduction in PPARGC1α and PPARα^57^. Together, these data indicate that short-term miR-424(322)/-503 overexpression disrupts lipolysis and fatty acid transport without directly impairing mitochondrial oxidative enzyme function.

**Figure 5.**
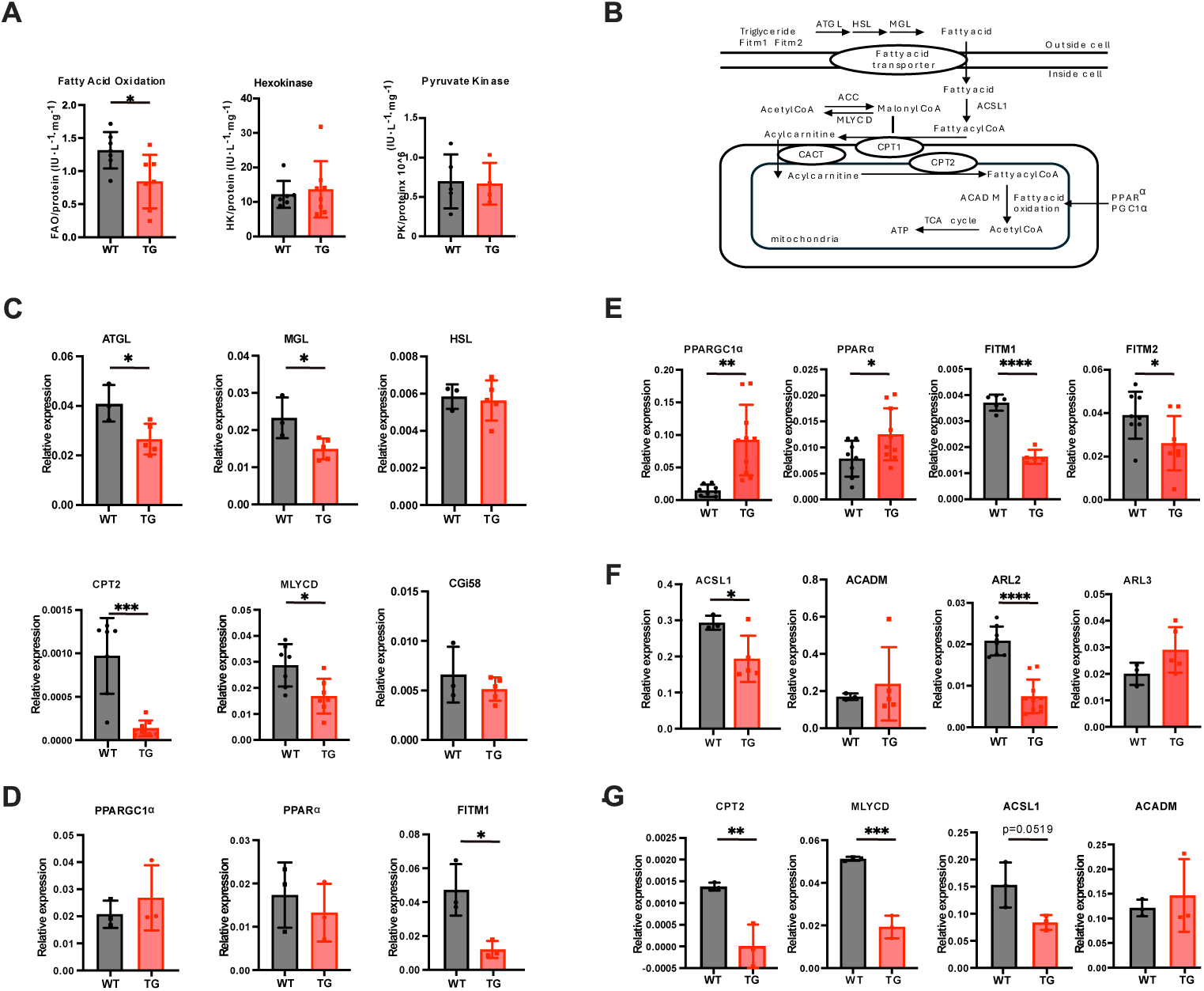
miR-424(322)/-503 overexpression suppresses cardiac fatty acid oxidation and lipid metabolic pathways. **(A)** Quantitative analysis of metabolic enzymatic activities, including fatty acid oxidation rate, hexokinase activity, and pyruvate kinase activity in WT and TG hearts following 7 days of miR-424(322)/-503 induction (WT n=5–7, TG n=4–7; Student’s *t*-test). **(B)** Schematic overview of cardiac lipid metabolic pathways highlighting key regulatory genes. **(C)** qPCR analysis of lipolytic genes adipose triglyceride lipase (ATGL), monoglyceride lipase (MGL), and hormone-sensitive lipase (HSL), comparative gene identification-58 (CGI-58), carnitine palmitoytransferase II (CPT2), malonyl-CoA decarboxylase (MLYCD), after 7 days of miR-424(322)/-503 induction (WT n=3, TG n=5; Student’s *t*-test). **(D)** qPCR analysis of transcriptional regulators peroxisome proliferator-activated receptor gamma coactivator-1α (PPARGC1α) and peroxisome proliferator-activated receptor-α (PPARα), as well as triglyceride storage proteins fat storage–inducing transmembrane protein 1 (FITM1) following short-term (2-day) miR-424(322)/-503 induction (WT n=5–8, TG n=7–8; Student’s *t*-test). **(E)** qPCR analysis of PPARGC1α, PPARα, FITM1 and FITM2 after 7 days of miR-424(322)/-503 induction (WT n=7, TG n=8; Student’s *t*-test). **(F)** qPCR analysis of acyl-CoA synthetase long-chain family member 1 (ACSL1) and acyl-CoA dehydrogenase medium chain (ACADM), along with ADP-ribosylation factor-like 2 (ARL2) and ADP-ribosylation factor-like 3 (ARL3) following 7 days of induction (WT n=3, TG n=5; Student’s *t*-test). **(G)** qPCR analysis of CPT2, MLYCD, ACSL1, and ACADM after short-term (2-day) miR-424(322)/-503 induction (WT n=3, TG n=3; Student’s *t*-test). *Date is represented as mean ± SEM. * p < 0.05*, *\*\* p < 0.01; *** p < 0.001 **** p < 0.0001*.

To further establish that these metabolic alterations represent primary early events, we analyzed gene expression after only 2 days of miR-424(322)/-503 induction. FITM1 expression was significantly reduced at this early timepoint, indicating rapid suppression of lipid storage. In contrast, PPARGC1α and PPARα expression remained unchanged, suggesting that compensatory transcriptional responses require longer induction (Figure 5D). Notably, MLYCD and CPT2 were already significantly downregulated by day 2, supporting the conclusion that impaired mitochondrial fatty acid transport is an early and direct effect of miR-424(322)/-503 overexpression. ACADM levels remained unchanged, and ACSL1 showed a downward trend that did not reach statistical significance (p = 0.0519) (Figure 5G). Collectively, these findings demonstrate that miR-424(322)/-503 rapidly impairs lipid storage and fatty acid transport as primary early events, preceding broader alterations in upstream regulatory pathways and mitochondrial oxidative enzymes.

## Discussion

Our findings demonstrate that H19X-encoded miR-424(322)/-503 plays a key role in both pathological remodeling and cardiac metabolism. Specific overexpression of the H19X-encoded miR-424(322)/-503 cluster in the heart leads to structural remodeling and dysregulation of cardiac function in a dose-dependent manner, and this deterioration is reversible upon withdrawal of the miR-424(322)/-503 overexpression. We also show that this microRNA cluster regulates cardiac metabolism, particularly fatty acid metabolism, during the early stages of functional disturbance, ultimately contributing to decline in heart function later on. Combined with our earlier work on the role of miR-424(322)/-503 in embryonic development, we propose a model for the function of miR-424(322)/-503 in cardiac development and disease. As some of the earliest non-coding RNAs expressed in cardiac progenitors, miR-424(322)/-503 contributes to cardiac lineage specification. Given its high expression during embryonic cardiac development and our finding that its re-expression in the adult heart suppresses fatty acid metabolism, we speculate that this cluster may also contribute to the relatively low reliance on fatty acid oxidation in the developing heart. Following birth, expression from the H19X locus declines, potentially releasing this inhibitory influence and facilitating the postnatal transition toward fatty acid oxidation as a major source of cardiac energy. In the adult heart, pathological reactivation of miR-424(322)/-503 may therefore partially recapitulate an embryonic-like metabolic program by attenuating fatty acid catabolism. However, the physiological role of miR-424(322)/-503 in developmental metabolic maturation and the contribution of its reactivation to cardiac pathophysiology remain to be determined.

These results align with previous clinical observations showing elevated H19X-encoded miR-424(322)/-503 in patients with heart failure^58,59^, as well as prior research demonstrating that overexpression of this cluster corelates with muscle atrophy^22,60,61^. Thus, the function of miR-424(322)/-503 illustrates the “fetal gene reprogramming” which manifests in adult cardiovascular and musculoskeletal diseases where stressed cells revert to an embryonic-like state regarding gene expression, contractility, and metabolism^62^. Classic fetal gene reprogramming involves shifts of contractile proteins from α-MHC to β-MHC^63^, re-expression of fetal isoforms of troponin T and troponin I^64^, increases in atrial natriuretic peptide and B-type natriuretic peptide^65,66^, and redeployed glucose transporter GLUT1 and key glycolytic enzymes. This metabolic reprogramming is rooted to the hypoxia environment in fetal hearts which rely on glycolysis rather than fatty acid oxidation for energy; stressed adult hearts adopt this reprogramming because they often experience hypoxia^67,68^. Therefore, this work adds a critical player in the fetal gene reprogramming paradigm. We have shown that the phenotypes are dose-dependent and can be reversed with the cessation of miR-424(322)/-503 induction, supporting that it is a potential target for alleviating heart failure.

Our data identify the miR-424(322)/-503 cluster as a major driver of cardiac dysfunction, characterized by rapid-progressing chamber dilation, thinning of the ventricular wall, extensive fibrosis, disorganized muscle fibers, and dysregulation of cardiac marker expression. This heart failure phenotype is among one of the strongest as compared to animal models where genes for sarcomere proteins, nuclear envelop proteins, cytoskeletal proteins were genetically manipulated^69–71^. Ablation of overexpression of miRNAs rarely produces such drastic phenotypes. miR-322(424) and miR-503 are members of the miR-15/16 family which are known to inhibit cell cycle progression^15,72^; while another member of the family, miR-195 was shown to regulate postanal cardiomyocyte cell cycle exit^73^. Overexpression of miR-195 caused hypertrophy which rapidly progressed to chamber dilation and systolic failure^74^; however, the rate and extent of cardiomyocyte deterioration was milder compared to miR-424(322)/-503 overexpression. Inhibition of miR-15 family by locked nucleic acid-modified anti-miR yielded protection against ischemia-reperfusion injury^75^. It is worth noting that miR-424(322)/-503 is highly specific for the myocardium, albeit the expression is restricted to embryos^17,22^. Other members of the miR-15/16 family have broad expression profiles including miR-195. The remarkable impact of miR-424(322)/-503 on fatty acid metabolism suggests that the function of miR-424(322)/-503 and other miR-15/16 family members may be more complex and propound than currently understood.

We observed no sex-dependent differences in the severity of cardiac dysfunction induced by miR-424(322)/-503 overexpression; both males and females exhibited comparable phenotypes. However, we anticipate that the pathophysiological function of miR-424(322)/-503 may be sex-related. The H19X locus is highly specifically expressed in female reproduction organs; this expression pattern is conserved in tetrapods^16^. It is expected that the expression from the H19X locus is subjected to hormonal regulation. Thus, although it is not within the scope of this study, it will be interesting to determine whether H19X ncRNAs are responsible for sex differences in cardiac energy metabolism in the future. It will be particularly interesting to investigate whether H19X non-coding RNAs play a role in the pathogenesis of female specific heart failure including peripartum cardiomyopathy and stress induced cardiomyopathy^76–78^.

Finally, we proposed that the miR-424(322)/-503 transgenic mouse is an inducible, versatile and reversible animal model that is useful in studying metabolic reprogramming in dilated cardiomyopathy and heart failure. One of the hallmarks of metabolic reprogramming is the shift of substrate utilization; downregulation of FAO is evident, accompanied by decreased expression of FAO-related genes^79^. However, ablation of the genes for key players in FAO, such as CD36, CPT1/2 and MLYCD, led to complex, context-dependent effects^80–85^, such genetic models are limited in studying metabolic reprogramming in dilated cardiomyopathy. The miR-424(322)/-503 transgenic mouse showed disruption of FAO with only two days of doxycycline induction, hypothetically due to multi-node regulation over FAO related genes. The disease course of this model is highly manipulatable, via changing the dose and frequency of doxycycline induction. We propose that this model will be useful for testing drugs as an intervention for progressive deterioration of energy efficiency. One of the weaknesses of this study is that we have not identified direct targets of miR-424(322)/-503 which can be used to explain the changes in metabolism and subsequent heart failure. Because the transcriptome of the TG animal enriched numerous predicted targets of miR-424(322)/-503, the phenotypes are likely mediated by a network of miR-424(322)/-503 target genes.

In summary, our study has three folds of significance for the understanding of cardiac metabolism and diseases. First, miR-424(322)/-503 is one of the rare regulators that are not a key enzyme or transporter of fatty acid but show a profound effect on fatty acid metabolism in the heart. Second, miR-424(322)/-503 may be targeted to reverse the course of heart failure if the role of it is established in humans. Finally, we have established a versatile animal model for heart failure. By altering the timing, frequency and dose of miR-424(322)/-503 induction, it will be possible to model heart failure at different stages and investigate cardiac metabolism as a key mechanism influencing heart function.

