## supplemental figures for "Overexpression of miR-424(322)/-503 Induces Severe Dilated Cardiomyopathy by Regulating The Fatty Acid Oxidation Gene Expression Program"

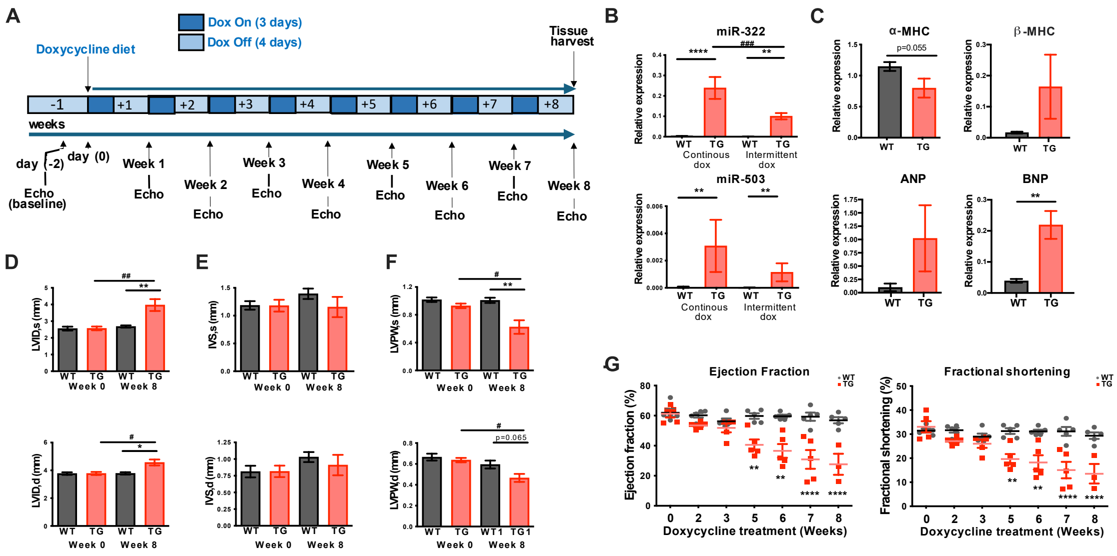

**Figure S1. Intermittent miR-424(322)/503 induction attenuates cardiac dysfunction and pathological remodeling.**

**(A)** Schematic timeline of intermittent doxycycline induction. Wildtype (WT) and Transgenic (TG) mice were placed on a cycling doxycycline (Dox) diet with 3 days "On" and 4 days "Off" per week for a total of 8 weeks. Echocardiography was performed to assess heart function weekly.

**(B)** qPCR analysis of miR-424(322) and miR-503 expression levels at week 8 following continuous Dox treatment and intermittent Dox treatment (WT n=6, TG n=5, student's t-test).

**(C)** qPCR analysis of cardiac hypertrophic markers α-MHC and β-MHC, and cardiac stress markers ANP and BNP following intermittent Dox treatment (WT n=6, TG n=5, student's t-test).

**(D-F)** Quantitative analysis of left ventricular internal diameter at systole (LVID;s) and diastole (LVID;d), left ventricular posterior wall thickness at systole (LVPW;s) and diastole (LVPW;d) and interventricular septal thickness at systole (IVS;s) and diastole (IVS;d) following intermittent Dox treatment (WT n=6, TG n=5, two-way ANOVA).

**(G)** Longitudinal echocardiographic assessment of left ventricular systolic function, including ejection fraction (EF, %) and fractional shortening (FS, %), measured at week 0, 2, 3, 5, 6, 7, and 8 following intermittent Dox treatment (WT n=5, TG n=5, two-way ANOVA).

*Date is represented as mean ± SEM. * $p < 0.05$, ** $p < 0.01$; *** $p < 0.001$ **** $p < 0.0001$*

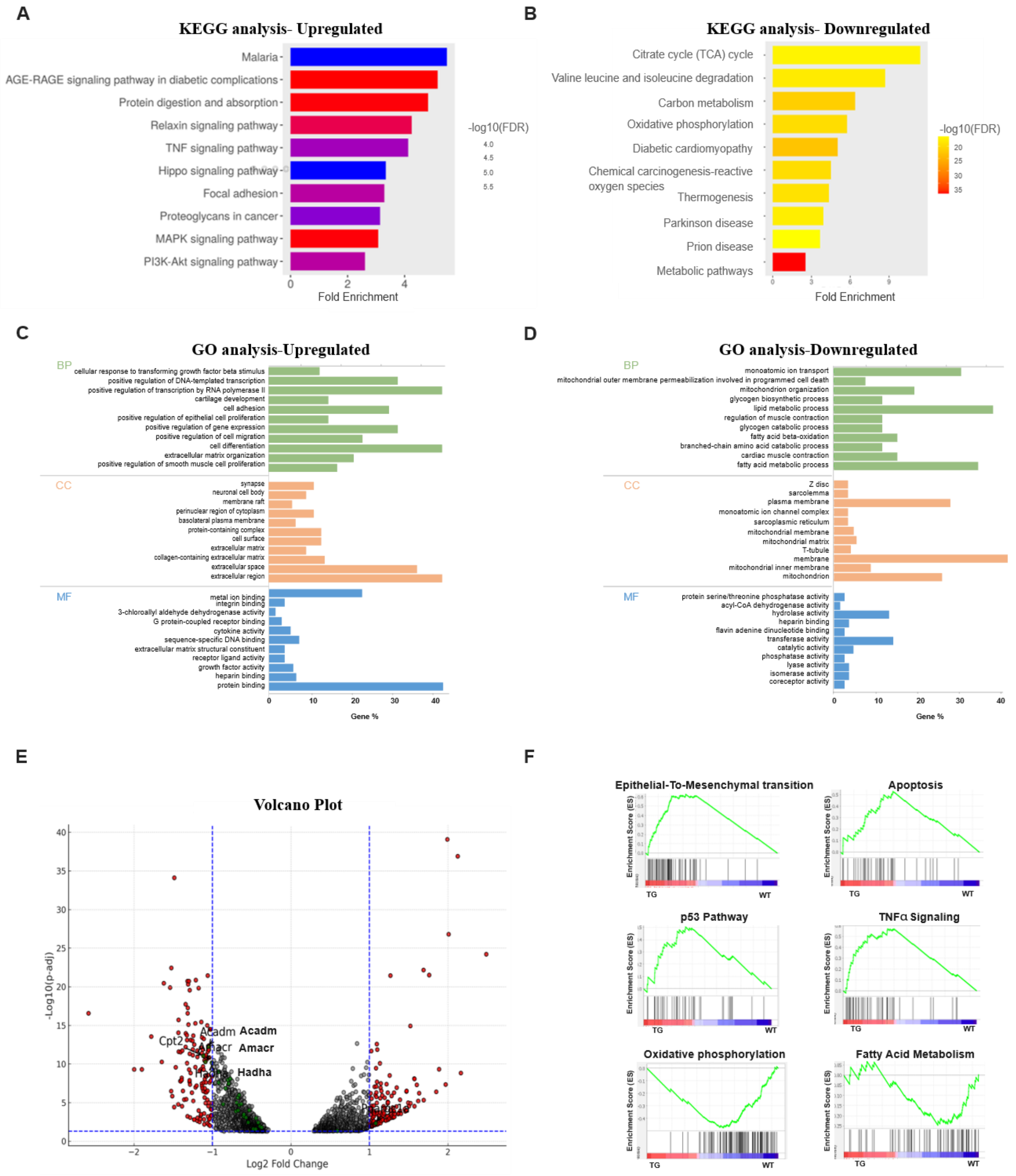


**Figure S2. Temporal transcriptomic profiling reveals a metabolic and signaling shift during the progression of miR-424(322)/503–induced heart failure.**

Transcriptomic profiling was performed on transgenic (TG) hearts at baseline (day 0) and after 14 days of miR-424(322)/503 induction.

**(A–B)** Kyoto Encyclopedia of Genes and Genomes (KEGG) pathway enrichment analysis identified significantly upregulated and downregulated pathways in TG hearts at day 14 compared with day 0.

**(C–D)** Gene Ontology (GO) enrichment analysis of the top 10 biological process (BP), cellular component (CC), and molecular function (MF) categories among upregulated and downregulated genes at day 14 relative to baseline.

**(E)** Volcano plot showing differentially expressed genes (DEGs) between day 14 and day 0, using a threshold of |$\log_2$ fold change| > 1. Fatty acid oxidation–related genes are highlighted in green, with genes exhibiting greater than twofold changes specifically labeled.

**(F)** Gene Set Enrichment Analysis (GSEA) of TG hearts at day 14 revealed enrichment of stress and remodeling pathways, including epithelial-to-mesenchymal transition, apoptosis, p53 signaling, and TNF-α signaling, alongside suppression of metabolic pathways such as oxidative phosphorylation and fatty acid metabolism.

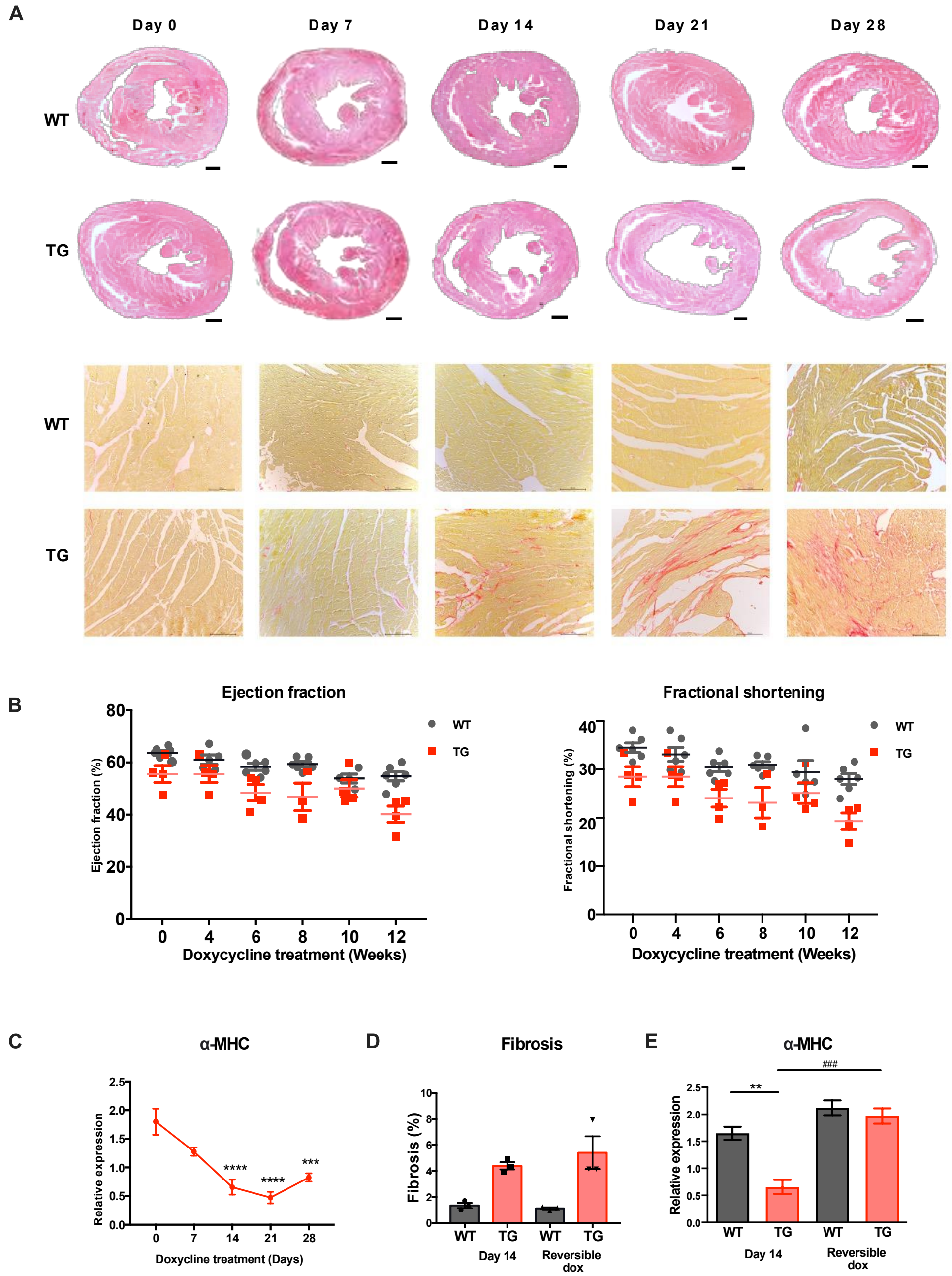


**Figure S3. Supplemental analysis of cardiac remodeling, dysfunction, and metabolic marker expression.**
**(A)** Representative hematoxylin and eosin (H&E) staining (upper panels; scale bar = 500 μm) and Picrosirius Red staining (lower panels; scale bar = 50 μm) of hearts from wild-type (WT) and transgenic (TG) mice at days 0, 7, 14, 21, and 28 of continuous doxycycline induction.
**(B)** Longitudinal echocardiographic assessment of left ventricular systolic function, including ejection fraction (EF, %) and fractional shortening (FS, %), measured at weeks 0, 4, 6, 8, 10, and 12 following low-dose doxycycline induction (200 mg/kg; WT n = 6, TG n = 5). Statistical analysis was performed using two-way ANOVA.
**(C)** Quantitative PCR (qPCR) analysis of the cardiac stress marker α-myosin heavy chain (α-MHC) expression in

WT and TG hearts at days 0, 7, 14, 21, and 28 of continuous doxycycline induction. Statistical analysis was performed using one-way ANOVA.
**(D)** Quantification of percentage change in Masson's trichrome–positive fibrosis following continuous doxycycline treatment compared with doxycycline withdrawal after 14 days of treatment in WT and TG mice (n = 3 per group).
**(E)** qPCR analysis of α-MHC expression following continuous doxycycline treatment compared with doxycycline withdrawal after 14 days of treatment in WT and TG mice (n = 3 per group).
*Date is represented as mean ± SEM. * $p < 0.05$, ** $p < 0.01$; *** $p < 0.001$ **** $p < 0.0001$*

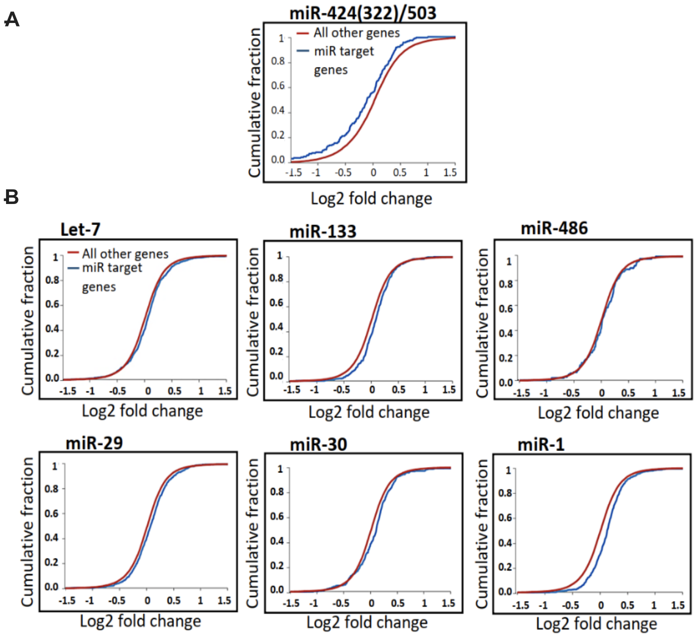


**Figure S4. Comparative transcriptomic impact of miR-424(322)/503 targets relative to other microRNAs.**
(A) Cumulative fraction (CF) plot comparing expression changes of predicted miR-424(322)/503 target genes (blue) versus non-target genes (red).
(B) CF plots comparing expression changes of predicted target genes (blue) versus non-target genes (red) for Let-7, miR-133, miR-486, miR-29, miR-30, and miR-1.
